# Global assessment of wildlife cryopreservation reveals major taxonomic gaps and strong species-specific differences in mammalian sperm cryotolerance

**DOI:** 10.64898/2026.05.22.727075

**Authors:** Gaukhar Y Iskakova, Alina Parkhomchuk, James K. Graham, Natasha S. Barteneva

**Affiliations:** Department of Biology, School of Sciences and Humanities, Nazarbayev University, 010000 Astana, Kazakhstan; Department of Biomedical Sciences, Colorado State University, 80523 Fort Collins, CO, USA; The Environmental Cluster, Nazarbayev University, 010000 Astana, Kazakhstan

**Keywords:** cryopreservation, wildlife, sperm, viability, motility, acrosome integrity, membrane integrity

## Abstract

Currently, global conservation efforts for wildlife focus on a limited number of cell types and species. Although protocols from domestic and non-threatened related species have been applied to endangered species, cryopreservation techniques are species-specific and are constrained by a lack of understanding of reproductive biology in these species. Based on a review of 126 original studies from 27 countries, encompassing 160 species, we assess the current state of cryopreservation in wildlife, including gametes, embryos, somatic cells, and various tissues. Furthermore, we focused on the most homogeneous and frequently studied cell type in wildlife cryobanking: mammals sperm.. A meta-analysis of 27 studies was conducted to examine species-specific and protocol-dependent factors that affect post-thaw sperm quality.

Our findings provide quantitative estimates of cryopreservation for various cell types and tissues in wildlife taxa. Furthermore, they serve as a crucial research roadmap, identifying major challenges in cryopreservation and proposing solutions.

## Introduction

Cryopreservation is a fundamental method for preserving a wide range of biological materials by cooling and storing them at ultra-low temperatures in liquid or vapor-phase nitrogen, effectively halting biochemical and metabolic activities and maintaining cellular viability for extended periods. The key challenges that cells must overcome at low temperatures are cell damage from intracellular ice crystal formation at high cooling rates, osmotic stress due to the solution effect at low cooling rates, and oxidative stress. To prevent the damaging effects of freezing, controlled rates and specialized cryoprotective agents (CPAs) are used, which affect the rates of water transport, nucleation, and ice crystal growth^1^. During warming and thawing, ice recrystallization can lead to cell swelling and potential rupture, while tissues and organs additionally experience non-uniform heating, inefficient warming rates, and uneven CPA redistribution^2^. Emerging technologies continue to evolve, including innovative CPA formulations, CPA-free methods^1,3^ , closed vitrification carriers^4^, and advanced warming techniques such as radiofrequency nanowarming^5^, along with microfluidics^6^, cryo-omics^7^, use of new cell types^8^ and artificial algorithms-enhanced methods^9–10^. Collectively, these advances require a new approach to integrate them into existing strategies, including assisted reproductive technologies (ART) and genome resource banking (GRB) for endangered wildlife.

Significant challenges in cryopreservation of cells and tissues from wild species are related to the limited availability of biological materials, small sample sizes, and poor integration of new methods. Successful handling, cryopreservation, cultivation, and subsequent use of cells and tissues require consideration of species-, cell-, and tissue-specific traits^11^. Protocols optimized for domestic livestock (e.g., bovine semen cryopreservation using glycerol-based extenders) frequently fail in nondomestic mammals due to differences in semen volume, pH, osmolarity, membrane permeability, and cellular resilience to cryoprotectants and cooling rates. Consequently, species-specific optimization—beginning with fundamental studies of gamete and tissue characteristics—is essential before scaling to assisted breeding. Among cells and tissues, oocyte cryopreservation is more complex mainly due to their large size, low surface-area-to-volume ratio, high water content, and complex cytoskeleton^12^. In contrast, sperm is the most extensively studied and most frequently banked cell type^13–14^. Overall, current wildlife cryopreservation research remains geographically and taxonomically skewed, is heavily centered on mammalian sperm and conventional freezing approaches, and consistently reports substantial post-thaw declines in key sperm quality parameters.

Here, we focus on the most recent global research in cryopreservation starting from 2020 to address two key research questions: 1. What are the sources of heterogeneity in wildlife cryopreservation results? 2. How to identify methodological and taxonomic gaps, and outline priority areas for future research and application in biodiversity conservation? We have combined a meta-analysis of wild species cryopreservation with a systematic review that covers different cryopreserved biological materials (gametes, somatic cells, and tissues), and preservation strategies in multiple taxonomic groups (mammals, birds, reptiles, amphibians, fish, and invertebrates), allowing us to consider cryopreservation in the context of inter-species and inter-tissue diversity, identify taxonomic and geographic patterns in the distribution of studies, and analyze the taxa-specific cryopreservation protocols used, including cryoprotectants, diluents, and freezing methods.

## Results

### Data Description

We extracted 126 studies from the initial 11,304 that met predefined selection criteria (Supplementary Table 1, 2, and Supplementary Figure S1). Although the initial search was limited to 2020, when analyzing systematic reviews, we rarely included earlier studies from 2018 to ensure the completeness and reliability of the final analysis. More than half of the recent studies on wild species cryopreservation come from just three countries: Brazil, Spain, and the United States. (Figure 1A).

**Figure 1.**
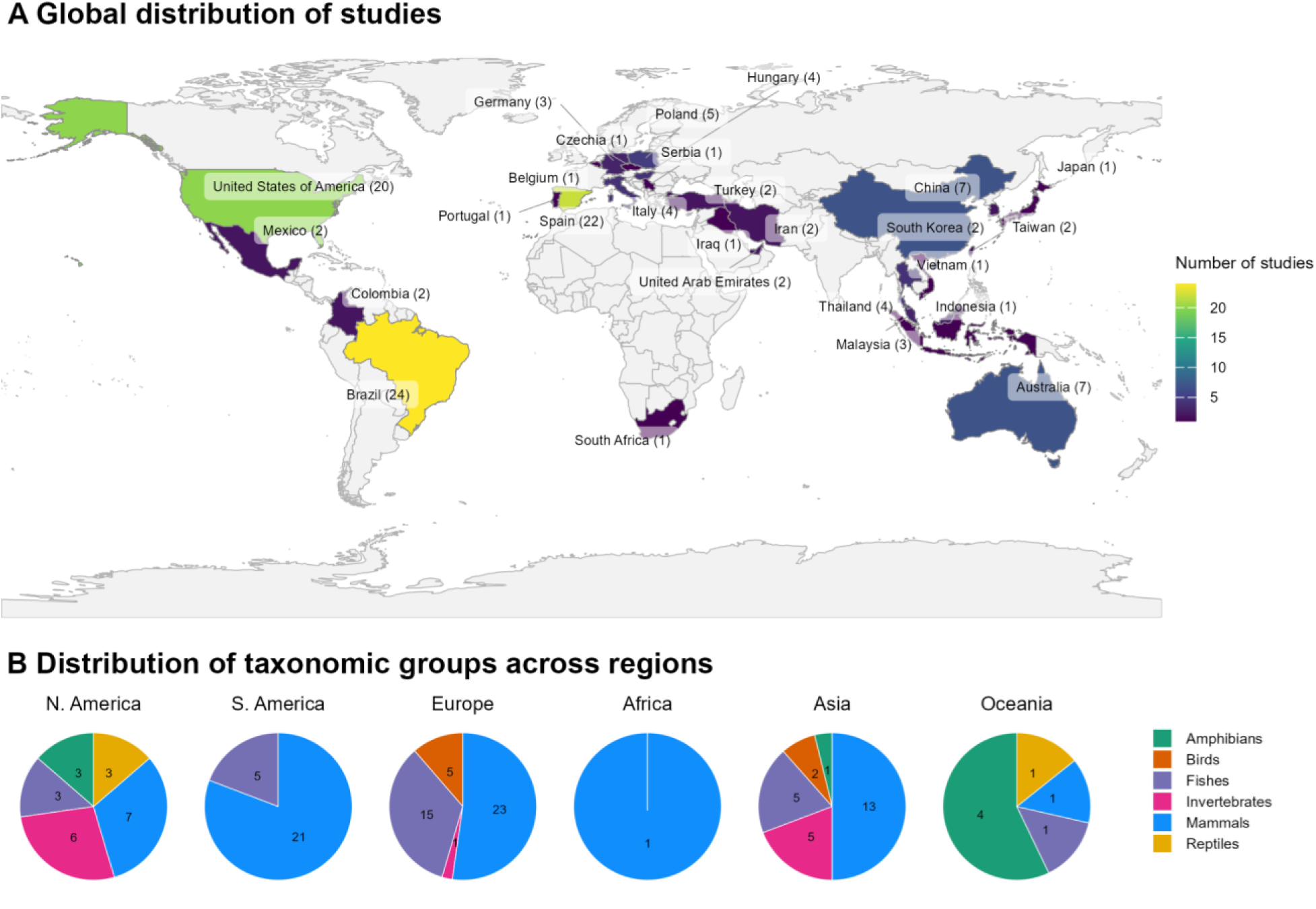
Global distribution of included studies by country and taxa. (**A**) The world map illustrates the geographic distribution of studies, with countries shaded according to whether they have at least one record. (**B**) Regional pie charts show the representation of taxonomic groups (amphibians, birds, fishes, invertebrates, mammals, and reptiles) in six geographic regions.

Furthermore, the articles were categorized by classes (Figure 1B), revealing that mammals (Mammalia) were the most common, followed by fish, amphibians, birds, and reptiles. Invertebrates were the subject of only 12 studies, but their representation was distributed across several classes, including Hexacorallia (7), Bivalvia (1), Cephalopoda (1), Gastropoda (1), Insecta (1), and Malacostraca (1). These results indicate a significant imbalance between the biological needs of species conservation and the current conservation efforts, as the most threatened taxa, such as amphibians and reptiles, are minimally represented.

Most studies reported cryopreservation of several, often closely related, species, encompassing 160 unique species, and resulting in 210 different records, and were primarily based on small cohorts of fewer than 10 individuals (Figure 2). Sample sizes varied significantly across studies. These included single-sample collections, such as postmortem extraction of testicular tissue or other biological materials, and protocols involving multiple-sample collections, such as repeated sperm sampling or fibroblast culture, which allowed cryopreservation of multiple samples from a single donor.

**Figure 2.**
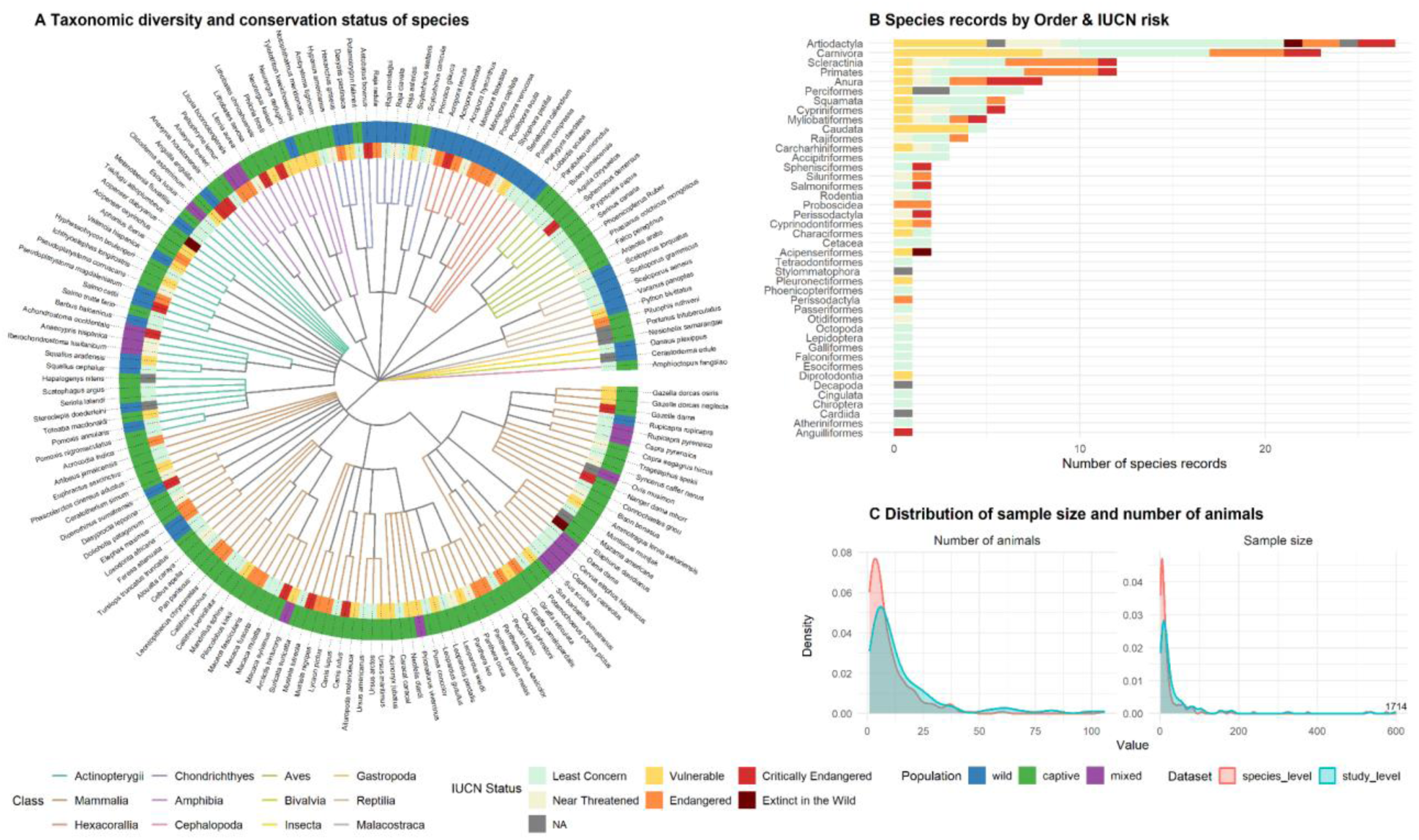
Taxonomic diversity and conservation status of the species included in this study. (**A**) The circle tree illustrates the taxonomic relationships among species, organized hierarchically from class to species level, with branches colored by taxonomic class. Two annotation rings provide conservation-related information for each species: the inner ring represents the International Union for Conservation of Nature (IUCN) Red List conservation status, and the outer ring displays the population status (wild, captive, or mixed). (**B**) The bar chart summarizes the number of species records by order, classified by IUCN conservation status, highlighting the distribution of extinction risk across taxonomic orders. (**C**) Density distributions show the number of animals and sample sizes at both the species (210 records) and study levels (126 studies).

Cryopreserved biological materials included gametes, gonadal tissue, and somatic cell lines. The choice of material and collection method reflected the species-specificities, logistical constraints, and conservation priorities. Preservation strategies varied: slow-freezing protocols were preferred for tissues, while vitrification was more common for highly sensitive cells such as mammalian oocytes or coral larvae. Sperm were the most frequently cryopreserved material among all taxa, accounting for 155 of the 210 experimental entries, and the most widely used methods were cryopreservation in liquid nitrogen vapor and slow programmable freezing. Pellet freezing was used primarily for certain mammals, birds, and reptiles. The disproportionate number of articles on sperm arises from the challenges associated with freezing female gametes and embryos.

Significant differences are observed in the methods used to obtain biological material, which vary depending on the anatomical and physiological characteristics of different species. For instance, abdominal massage was used in ray-finned fishes, cartilaginous fishes, birds, and reptiles. In non-mammals cloacal massage was observed in studies involving amphibians and birds, while spermic urine collection was predominant in amphibian. Additionally, gamete bundles were collected in cases involving stony corals. In Hexacorallia, samples were obtained by collecting colony fragments; and in one instance, thermal shock was applied to stimulate gamete release in Bivalvia. In Actinopterygii, gonad dissection was used to extract testes, ovaries, spermatogonial cells, and primordial germ cells (PGCs), while ovarian tissue excision was used to extract ovarian tissue and stem cells. In gastropods, ovotestis dissection was used. On the other side, in mammals, ejaculated spermatozoa were primarily obtained by rectal probe electroejaculation (EE) or transrectal ultrasound-guided massage of the accessory sex glands (TUMASG). Less commonly used methods included urethral catheterization (UC), penile vibrostimulation (PVS), penile electrostimulation (PES), among others (Figure 3A).

**Figure 3.**
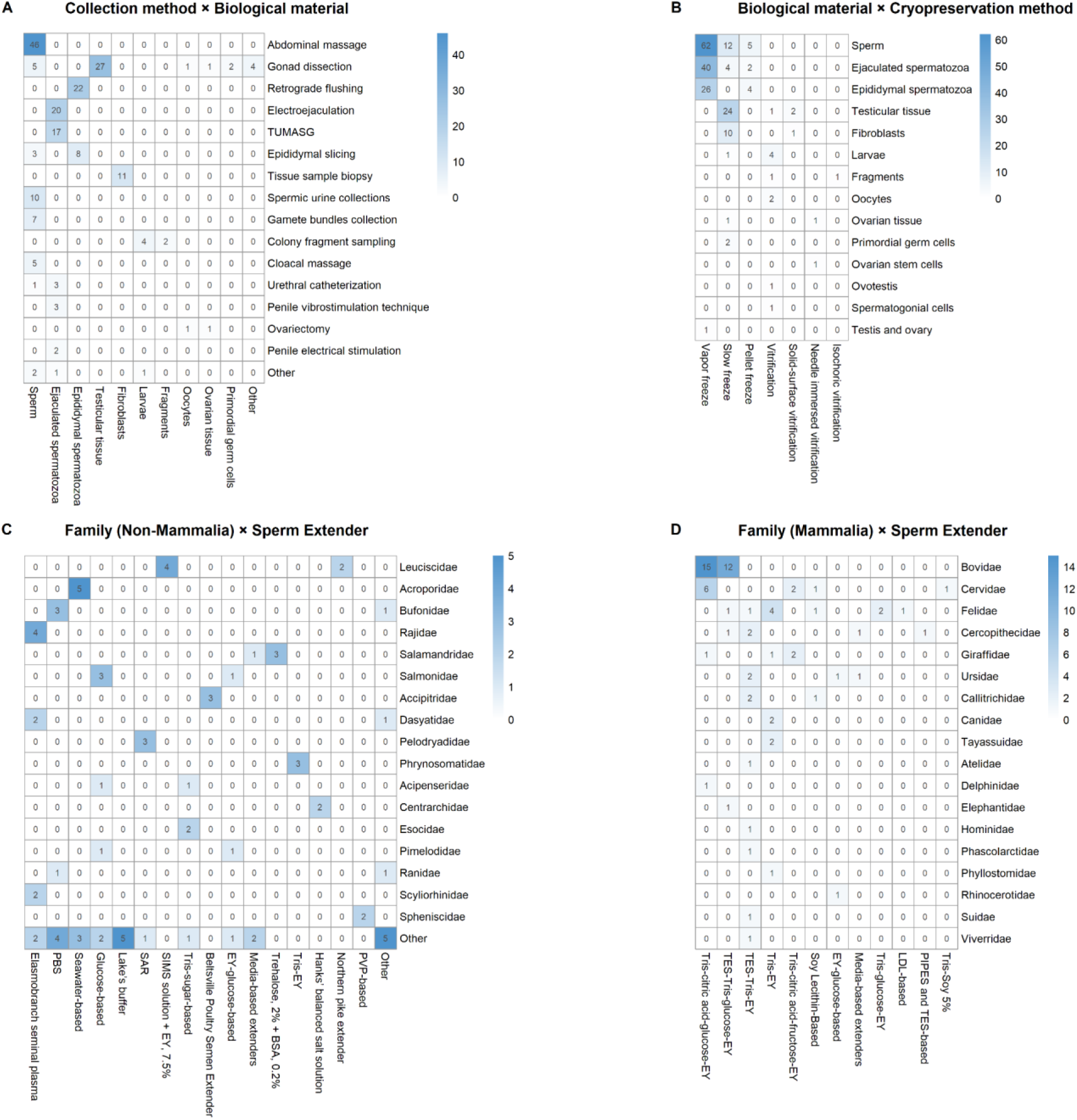
Multi-panel heatmaps showing relationships between **(A)** collection methods and biological materials, **(B)** biological materials and cryopreservation methods, **(C)** extenders and families (non-Mammalia), and **(D)** extenders and families (Mammalia). Color intensity corresponds to the number of observations.

In fishes, gonadal tissue and primordial germ cells from Actinopterygii were cryopreserved using slow programmed freezing; ovarian stem cells were preserved using needle-immersed vitrification (NIV), while spermatogonial cells were also vitrified. In invertebrates, Gastropoda ovotestes and Hexacorallia larvae and fragments were vitrified, including one case of isochoric vitrification.

Mammalian testicular tissue was typically cryopreserved using slow freezing or solid-surface vitrification (SSV). In studies involving fibroblasts, seven research groups employed slow programmable freezing, and one reported that SSV was as effective as slow freezing in preserving the skin of six-banded armadillo^15^. Oocytes have been vitrified, and ovarian tissue has been preserved using NIV. It is important to note that several papers compare different cryopreservation methods. Based on their experimental results, we included the protocol that provides the best survival and post-thaw outcomes (Figure 3B).

The combinations of cryoprotective agents (CPAs) and extenders, as well as their concentrations, varied significantly across cryopreservation methods, tissue types, and animal species involved. Details regarding the combinations of sperm extenders are presented in Figures 3C and 3D, while the sequential workflow—including the order, biological material, collection method, and CPAs—is depicted in Figure 4.

**Figure 4.**
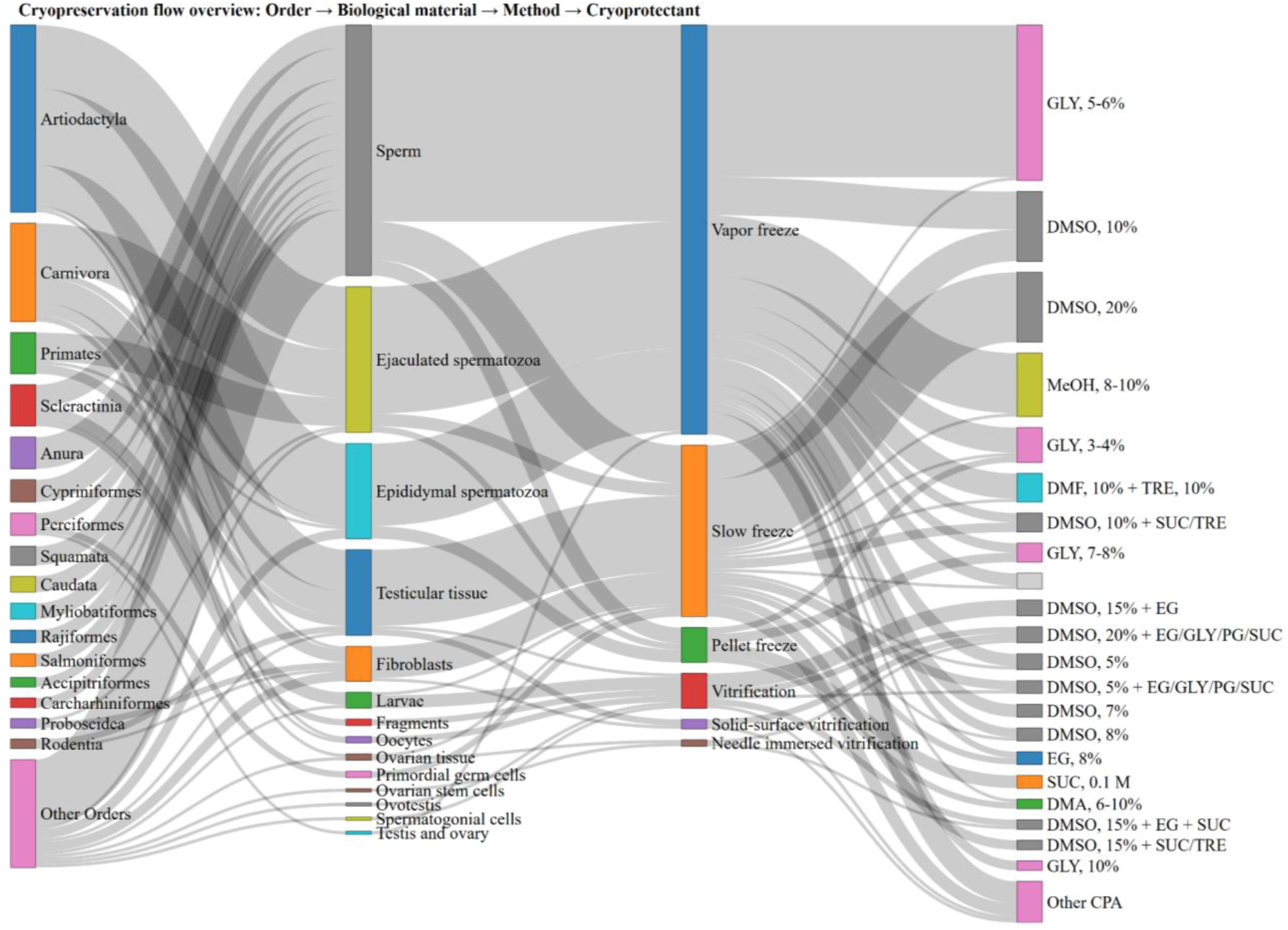
Sankey diagram illustrating the flow of cryopreservation methods across taxonomic orders, biological materials, cryopreservation methods, and cryoprotective agents in the analyzed dataset. Node height is proportional to the number of samples. Rare taxonomic orders and cryoprotective agents (n < 3) were grouped into the categories "Other orders" and "Other CPA," respectively. Biological materials and cryopreservation methods are shown without grouping. The diagram demonstrates the dominant combinations and pathways in cryobanking practices.

Tris-based extenders containing egg yolk were the most commonly used formulations for cryopreservation of mammalian sperm (Figure 3D). This prevalence likely stems from the widespread adoption of protocols optimized for bull semen, in which Tris serves as a simple, inexpensive buffer that maintains a pH of ∼7.0, and egg yolk (EY, or milk-based components) protects against cold shock during cooling. In contrast, non-mammalian sperm (particularly amphibians and fish) are often frozen without egg yolk (Figure 3C), reflecting higher tolerance to rapid cooling and differences in membrane lipid properties. These patterns highlight both historical reliance on established protocols and taxon-specific differences in sperm cold-shock sensitivity.

The significant taxonomic and methodological diversity observed in wildlife cryopreservation studies is evident from the various approaches taken for different species. Protocols are often tailored to the specific biological constraints and practical considerations unique to each species. While this adaptability reflects the versatility of cryopreservation methods, it also leads to significant variability in experimental design, outcome measures, and reporting practices. This variability complicates direct quantitative comparisons across taxa and tissue types. In contrast, mammalian sperm cryopreservation represents the most extensively studied and methodologically consistent subset, characterized by standardized outcome measures. This uniformity facilitates robust quantitative synthesis and systematic evaluation of factors influencing post-thaw sperm quality, providing a solid foundation for meta-analysis.

### Pooled analysis

For the meta-analysis, data on sperm parameters were pooled from 27 experiments (23 studies, Supplementary Tables 3, 4, and 5), including total motility, progressive motility, viability, normal morphology, acrosome integrity, membrane integrity, and mitochondrial activity. Heterogeneity was extremely high (I² > 90%) for all parameters, indicating significant between-study variability, which predefined a choice of a random-effects model.

As expected, the meta-analysis using a random-effects model revealed a decrease in all sperm parameters after cryopreservation (Figure 5). Separate scatter plots for each parameter are available in Supplementary Figures S2–S8.

**Figure 5.**
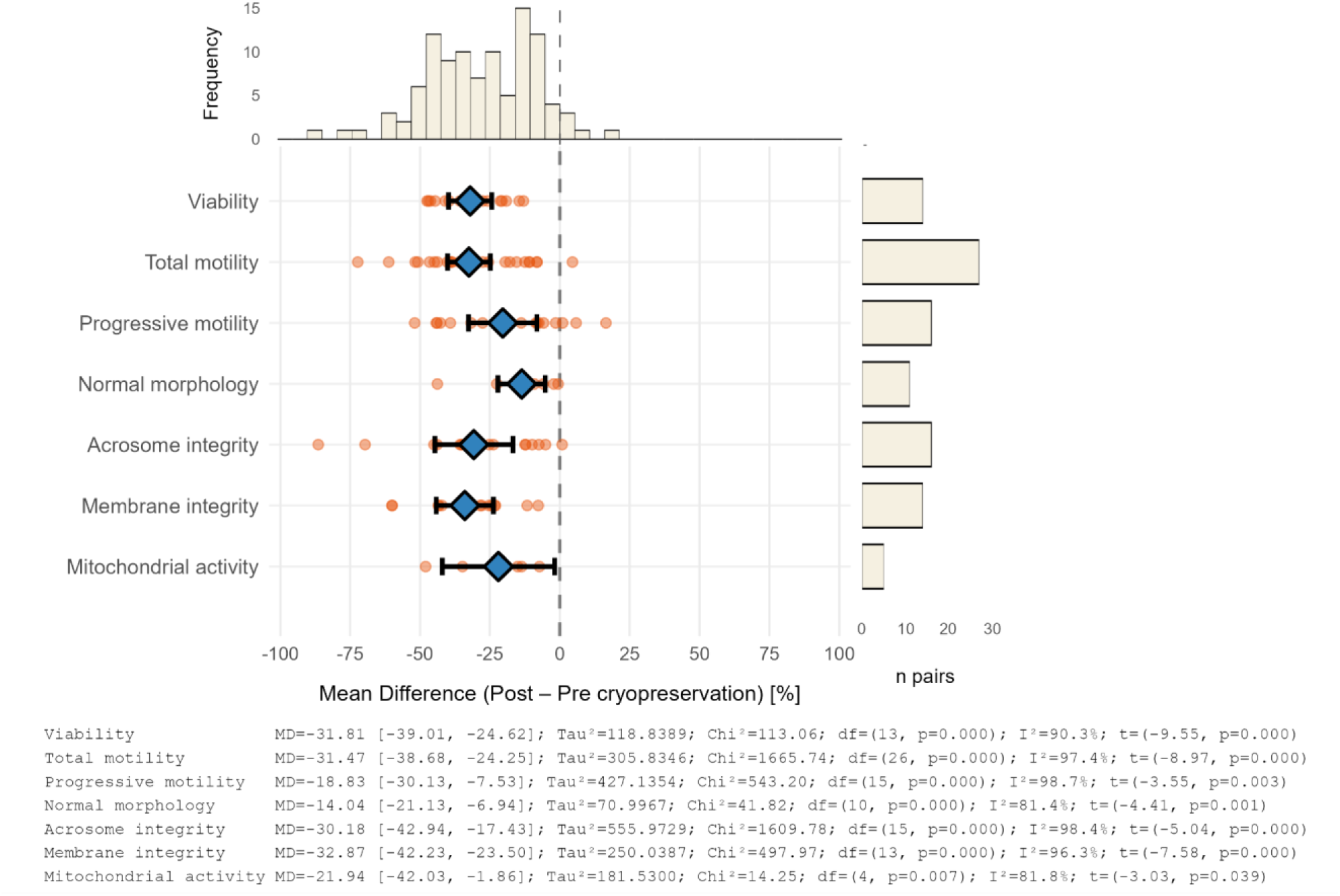
The effect of cryopreservation on sperm parameters: viability, total motility, progressive motility, normal morphology, acrosome integrity, membrane integrity, and mitochondrial activity. The central forest plot displays the effect sizes of individual studies (● - orange dots) with 95% confidence intervals (— horizontal lines) and the pooled effect estimates (♦ - blue diamonds) calculated using a random-effects multilevel model (REML with Knapp-Hartung correction). The histogram above the plot shows the distribution of effects for all outcomes. The bar plot on the right indicates the number of paired comparisons (post-pre-cryopreservation) for each outcome. The bottom panel presents full statistics for each outcome, including MD with 95% CI, heterogeneity estimates (τ², I²), Chi-square test for heterogeneity (Q, df, p), and the overall effect (t, p).

The morphology of sperm was the least affected by the freezing and thawing process, showing only a 14% reduction in normal morphological forms. Progressive motility decreased by 19%, while mitochondrial activity dropped by 22%. Other parameters, including viability, total motility, acrosomal and membrane integrity, decreased by approximately 30%. These changes reflect the combined effects of cold stress, osmotic changes, and the toxic effects of cryoprotectants.

### Sensitivity Analyses

The sensitivity analysis showed that the pooled mean differences (MDs) remained mostly stable across all outcomes. Removing outliers and influential studies had minimal impact on viability, motility, morphology, and other metrics. Heterogeneity was significantly reduced after removing outliers for total motility and normal morphology, and to a lesser extent for other parameters, suggesting that few studies accounted for most of the variability observed in the original analysis. All diagnostic influence plots, such as the Baujat plot, leave-one-out plot, Cook’s distance, etc., are presented in Supplementary Figures S9–S24.

To explore possible sources of heterogeneity in post-thaw semen parameters, subgroup analyses were conducted. Except total motility, other semen parameters were reported inconsistently across studies and either lacked sufficient data for meaningful subgroup analyses or did not demonstrate statistically significant subgroup effects.

Post-thaw sperm viability was significantly affected by population origin (captive versus wild), sperm collection method, and the cryoprotective agents (*p* = 0.0360, *p* = 0.0016, *p* = 0.0361). No significant subgroup effects were observed for factors such as taxonomic family, extinction risk, breeding season, collection frequency, biological material (ejaculated versus epididymal), or extender composition. Total motility demonstrated significant differences between taxonomic family, extinction risk, collection method, cryoprotectant agent, and extender type. In contrast, population origin, breeding season, collection frequency, and biological material did not show significant effects. Similarly, progressive motility was notably influenced by population origin, collection method, cryoprotectant agent, and extender (all *p* < 0.0001). Acrosome integrity did not differ across any of the examined subgroups. For membrane integrity, only the cryoprotective agent exhibited a statistically significant effect, and for normal morphology only the type of extender, with all other subgroup variables remaining non-significant.

The method of sperm collection has a significant impact on the quality and survival of the sperm. In published studies, EE and TUMASG have been two most commonly used techniques for collecting semen from wild mammals. A less stressful method is TUMASG, but the effectiveness of this technique is species-specific, with the greatest success observed in ruminants^16^. Epididymal sperm extraction methods involve collecting sperm directly from the epididymis after euthanasia, castration, or accidental death. This approach is limited by the fact that it does not allow for a second sampling^17^.

It is also important to consider the composition of extenders, including the CPA (usually glycerol), the buffer utilized (Tris, TES, HEPES, etc.), and the source of low-density lipoproteins (usually egg yolk or milk) needed to prevent ‘cold shock’ for sperm that are sensitive to this issue. When evaluating glycerol concentrations, lower levels (less than 5%) were associated with significant declines in overall motility, progressive motility, and membrane integrity. The lowest reduction in sperm quality was noted at a concentration of 5% glycerol. Other cryoprotectants and higher glycerol concentrations (greater than 5%, with data available only for total motility) yielded intermediate outcomes. However, concentrations exceeding 6-8% can be toxic, negatively affecting sperm motility, fertility, and membrane integrity.

In addition, the cooling rate and other factors also influence post-thaw sperm survival. Among extenders, TCGey (Tris, citric acid, glucose, and egg yolk) had the least effect on overall motility, progressive motility, normal morphology, and acrosomal integrity, but it should be noted that the choice of extender is highly species-specific. Since sperm from some species are not sensitive to ‘cold shock’ and therefore do not need egg yolk, in other instances, the sperm may prefer a higher pH, and a different buffer that maintains the desired pH more efficiently may be used.

The dominance of one technique in the available datasets made it challenging to compare different cryopreservation methods. Total motility showed no significant difference between slow freezing and vapor freezing. Acrosome integrity was found to be better with slow freezing compared to vapor freezing, while membrane integrity was superior with vapor freezing than with other methods, although no statistically significant differences were found.

### Publication Bias

The visual analysis of funnel plots indicated no evidence of publication bias (Supplementary Fig. S25-27). Egger’s regression test also showed no signs of funnel plot asymmetry (all *p* > 0.05, Supplementary Table 7). The trim-and-fill method did not detect any missing studies in any parameters except AI, which had three potentially missing studies on the left side of the funnel plot. The estimate of the corrected effect remained close to the original pooled effect.

## Discussion

Cryopreservation has evolved from a specialized laboratory procedure into a fundamental component of modern biological research, biotechnology, and conservation^12, 18–19^.

At the cellular level, cryopreservation involves dynamic volume changes that occur during the introduction and removal of cryoprotective agents, dehydration during freezing, and rehydration during thawing and dilution (Figure 7).

**Figure 6.**
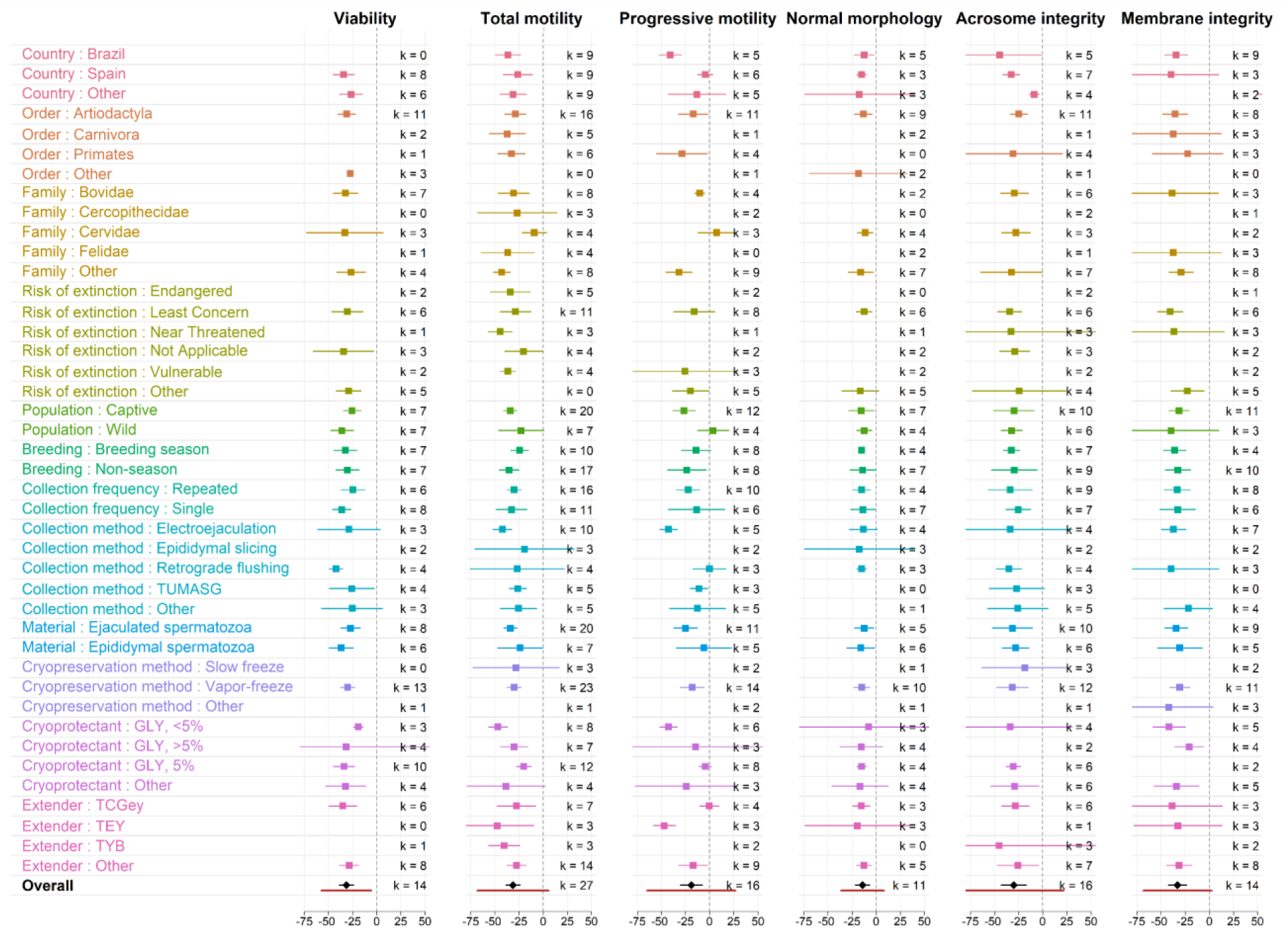
Grouped forest plot shows effect sizes grouped by category (viability, total and progressive motility, normal morphology, acrosome and membrane integrity). A vertical dashed line at zero indicates no effect. The square (◼) represents the effect of the individual subgroup, the horizontal line (—) represents the confidence interval, the black diamond (♦) represents the overall effect, the red horizontal line (—) represents the prediction interval, and *k* is the number of pairwise comparisons for each variable.

**Figure 7.**
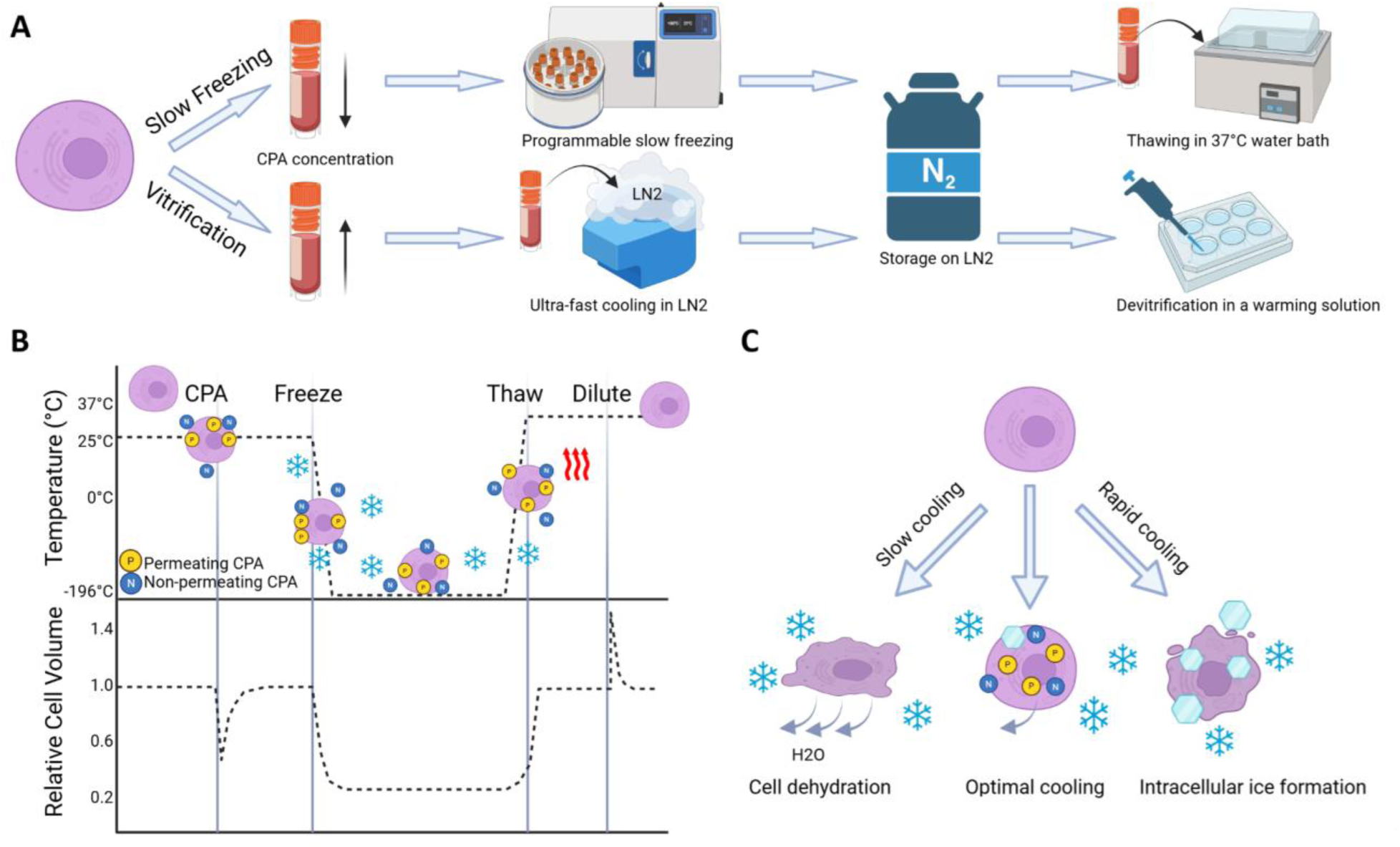
General principles of cryopreservation and cellular responses to cooling and thawing. (**A**) Schematic representation of a typical cryopreservation protocol. (**B**) Typical temperature profile and associated cell volume changes during cryopreservation. Adapted from^20^.(**C**) Mazur’s two-factor hypothesis. An optimal, cell-type-specific cooling rate minimizes both dehydration damage and intracellular ice formation, thereby maximizing survival after thawing. Adapted from Mazur et al. (1972)^21^.

When sperm are cooled to 5°C, the membranes of most sperm undergo a phase transition from the fluid to a gel-like state. This change can damage the membranes, causing the cells to undergo a phenomenon known as ‘cold shock’, which compromises the integrity of the in sperm cells. The susceptibility to this phase change is dependent upon the lipid composition (yolk in non-mammals) of the sperm. Sperm with higher cholesterol to phospholipid ratios, such as human and rabbit sperm ^22^, or those with elevated levels of unsaturated lipids, like chicken sperm^23^, are more resilient to cooling than sperm from species that are typically more vulnerable, such as bull, ram, stallion. Sperm from species that are susceptible to cold shock can be protected from damage by adding low-density lipoproteins (LDL) to the sperm prior to cooling^20^. LDL are found in high concentrations in egg yolk and in milk, which is why most sperm extenders used for cooled semen or cryopreservation contain either egg yolk or milk.

The cholesterol content of sperm can be increased using cholesterol-loaded cyclodextrins (CLC), and after CLC treatment, sperm from species that are cold shock sensitive survive cooling^24–25^ and cryopreservation^24^ more effectively.

The addition of and particularly the removal of CPAs affects cell volume (Figure 7B), which can damage sperm. Since water is more permeable to cell membranes than CPA, when CPAs are added to cells, the cell initially shrinks and then regains its volume as the CPA slowly equilibrates into the cell. On the other hand, when a cryopreserved cell loaded with CPA is diluted into a CPA-free medium, water rushes into the cell, causing swelling before the CPA can slowly leave the cell. The osmotic tolerance of a cell evaluates how much that cell can expand without damage^26^. If a cell swells beyond its capacity, it will rupture causing cell death, and cells with a narrow osmotic tolerance can benefit from using CPAs that are more permeable to the cell membrane^27^. Although glycerol is the most commonly used CPA to cryopreserve sperm, using CPAs with higher cell membrane permeability can improve the cryosurvival rate of sperm from several domestic^27–28^ and wild animals^29–30^.

There is often difficulty in collecting sufficient sample numbers or sample volumes from wild species, and the difficulty and expense of determining either the lipid composition or osmotic tolerance of these sperm. Therefore, several studies have been done in which sperm from wild species were treated with CLC prior to freezing (monkey^31^; carp^32^, camel^33^, gazelle^34^) or were frozen using alternative CPAs (bat and fish^30,35–38^, without knowing either the lipid composition or the osmotic tolerance of the sperm.

### Mammals

Most of our knowledge of preserving genetic material is based on the success of cryobanking and ART in humans, livestock, and laboratory animals, and these methods are easily adapted to wild mammals. However, most attempts at cryobanking in wild animals remain limited to a few species, primarily mammals, and biological material types, primarily spermatozoa. Currently, the true integration of ART into population management has been achieved only in a few species. Notable examples include the black-footed ferret (*Mustela nigripes*) in the USA and the giant panda (*Ailuropoda melanoleuca*) in China, where artificial insemination using thawed semen has become a routine and effective component of conservation programs^39–40^ .

Similar progress is being made in the conservation of white rhinoceroses. Advances in artificial insemination have led to the successful birth of multiple southern white rhino calves from fresh, chilled, or frozen-thawed semen^41^. Additionally, complementary approaches, such as autologous grafting of cryopreserved prepubertal testicular tissue, have successfully restored spermatogenesis and generated functional sperm capable of producing healthy offspring in rhesus macaques^42^, highlighting the broader potential of gonadal tissue banking and transplantation for fertility rescue. Less research has been devoted to preserving female genetic material, as the task is more complex in the case of oocytes, and obtaining the material is challenging^43–44,12^. Oocytes have relatively low permeability to cryoprotectants and water due to a thick outer cell membrane, which increases the risk of ice crystal formation^45^. In some species, an additional challenge is high lipid content in the cytoplasm. This prompts the exploration of alternative strategies, such as gonadal tissue xenotransplantation, *in vitro* culture systems, and interspecies somatic cell nuclear transplantation.

Testicular tissue contains a large number of early-stage germ cells that can mature *in vitro* or through grafting, making them useful for ART. Typically obtained postmortem, testicular tissue fragments can be cryopreserved using various methods; however, there is significant variability in tissue cryosensitivity. For example, feline testicular tissue better survives vitrification, while rodent and artiodactyl tissues tend to withstand slow freezing more effectively. Nonhuman primates show promising results with both methods^46–47^. Similarly, ovarian tissue can be frozen either by slow freezing or vitrification^47^ .

Cryopreservation of somatic and stem cells is less complex than that of germ cells. However, using these cells for cloning and the production of induced pluripotent stem cells (iPSCs) is difficult; low live birth rates, low neonatal survival rates, and developmental abnormalities among clones have been documented in the literature^48^. However, somatic cell nuclear transfer (SCNT) technology, using cell lines from tissues frozen 30–40 years ago, has been able to achieve the birth of clones of the footed ferret (*Mustela nigripes*) and the Przewalski’s wild horse (*Equus ferus przewalskii*). This advancement has the potential to diversify the genome and increase the size of these populations^48^.

### Birds

Sperm freezing in avians is complicated by the shape of spermatozoa and their membrane permeability. Standard glycerol has a pronounced contraceptive effect in birds, requiring additional steps to remove it before sperm use^49^ or use alternative CPAs^50^ . Successful examples of wild bird sperm cryopreservation include cranes (*Grus americana, Grus vipio*), penguins (*Spheniscus demersus, Pygoscelis papua*), and Arabian Bustard (*Ardeotis arabs*)^16,51–52^. In Japanese quail (*Coturnix japonica*), testicular tissue retains viability and spermatogenic function after transplantation into castrated recipients. Intramagnal insemination of fluid from these transplants produces live donor-derived offspring, demonstrating functional recovery^53^.

The freezing of mature oocytes is currently not possible in fishes, amphibians, reptiles, and birds^11^. However, ovarian cells can be used to preserve female genetic material. After vitrification, these cells can be recovered through transplantation, leading to the production of donor-derived chicks. The use of primordial germ cells for cryopreservation and transplantation into surrogate hosts has been documented in chickens^54^. An alternative to germ cells is the cryopreservation of follicle cells, which can be obtained by plucking feathers. The isolation of somatic cells and, potentially, stem cells from the follicle holds promise for biobanking and the future production of clones.

### Reptiles

Reptiles represent a diverse and species-rich group known for their complex reproductive systems. These systems are different in oviparous and viviparous species, in species with large yolk-bearing eggs, long-term sperm storage in the female tract, a complex genetic and temperature-dependent sex determination system, sex reversal, and many other features. Protocols have been published for lizards and squamates, but research primarily focuses on sperm cryopreservation, and the overall number of studies remains low^55–56^ . Notably, Sandfoss and co-authors^57^ reported the first successful case of artificial insemination of Louisiana pinesnakes (*Pituophis ruthveni*) using thawed semen, resulting in the birth of three viable hatchlings.

### Amphibians

Amphibian sperm can remain viable for weeks at low temperatures and are highly tolerant of exposure to cryoprotectants. While there are many examples of the use of stored sperm and artificial fertilization leading to the production of live offspring, only a limited amount of species have been studied^58^. For example, among endangered species, thawed sperm has been used in ART for *Pseudophryne corroboree*^59^, *Rana sevosa*^60^, *Anaxyrus boreas boreas, Lithobates sevosa* and *Anaxyrus fowleri*^61^, *Cryptobranchus alleganiensis*^62^, and *Peltopryne lemur*^63^. Cryopreserved sperm has produced viable F1 and even F2 offspring in several species, with some individuals reaching sexual maturity and contributing to reintroduction efforts.

To preserve female fertility, the cryopreservation of undifferentiated primordial germ cells may be the only option. While cryopreservation of somatic tissues shows promise, one of the major challenges in freezing amphibian skin and cell cultures is the contamination from bacteria and fungi, which can be difficult to eliminate^64^.

### Fishes

Sperm cryopreservation has been successfully applied to many commercial fish species, mostly salmonids, cyprinids, and sturgeons, but remains poorly studied in wild species. Significant differences in sperm cryosensitivity and the lack of species-specific protocols limit wider application in conservation. Currently, there is no effective method for cryopreserving the maternal genome, making the use of PGCs, spermatogonia, and oogonia, as well as germ cell transplantation, the most promising for restoring endangered species and genetic diversity^65^. Additionally, somatic cells, as well as skin and fin tissues in fish, are interesting options for genome preservation as they carry both the maternal and paternal genomes^66^.

### Invertebrates

Invertebrates are often overlooked in the cryopreservation literature, despite being the most species-rich class. For many species, no protocols have yet been developed, except for certain corals and some insects. Currently, asexual propagation and colony fragmentation methods are primarily used; however, these methods only yield clonal material. To maintain genetic diversity sexual reproduction is preferable. There are successful cases of sperm cryopreservation utilizing a standardized cryopreservation protocol for over 30 different coral species^67^. Recently, coral larvae were cryopreserved using vitrification and an advanced laser-warming process, which avoided damage due to ice formation^68^. However, effective cryopreservation technologies for other cell types are not yet sufficiently developed for reef restoration programs^69^.

Research on cryopreservation in insects has focused on model organisms such as silkworms and fruit flies, as well as agriculturally important honeybees. Recently, several studies and protocols for the cryopreservation of *Anopheles gambiae* and *Anopheles stephensi* were published, which may help in the cryopreservation of other invertebrates with early development stages in aquatic environments^70–71^.

### Persistent Challenges and Future Trajectories in Wildlife Cryopreservation

Over the past two decades, progress has been made in the development and application of germ cells cryopreservation and *in vitro* technologies. However, significant challenges remain, including the need for research to understand the biological characteristics of different species, improve post-thaw results, and establish standardized and reproducible protocols for a given species. A well-established infrastructure framework is required, encompassing ethical and legal collection, transfer to biobanks/biorepositories, standardized processing and quality control, and secure long-term storage (Figure 8). Such infrastructure is often lacking in biodiversity hotspots where it is needed.

**Figure 8.**
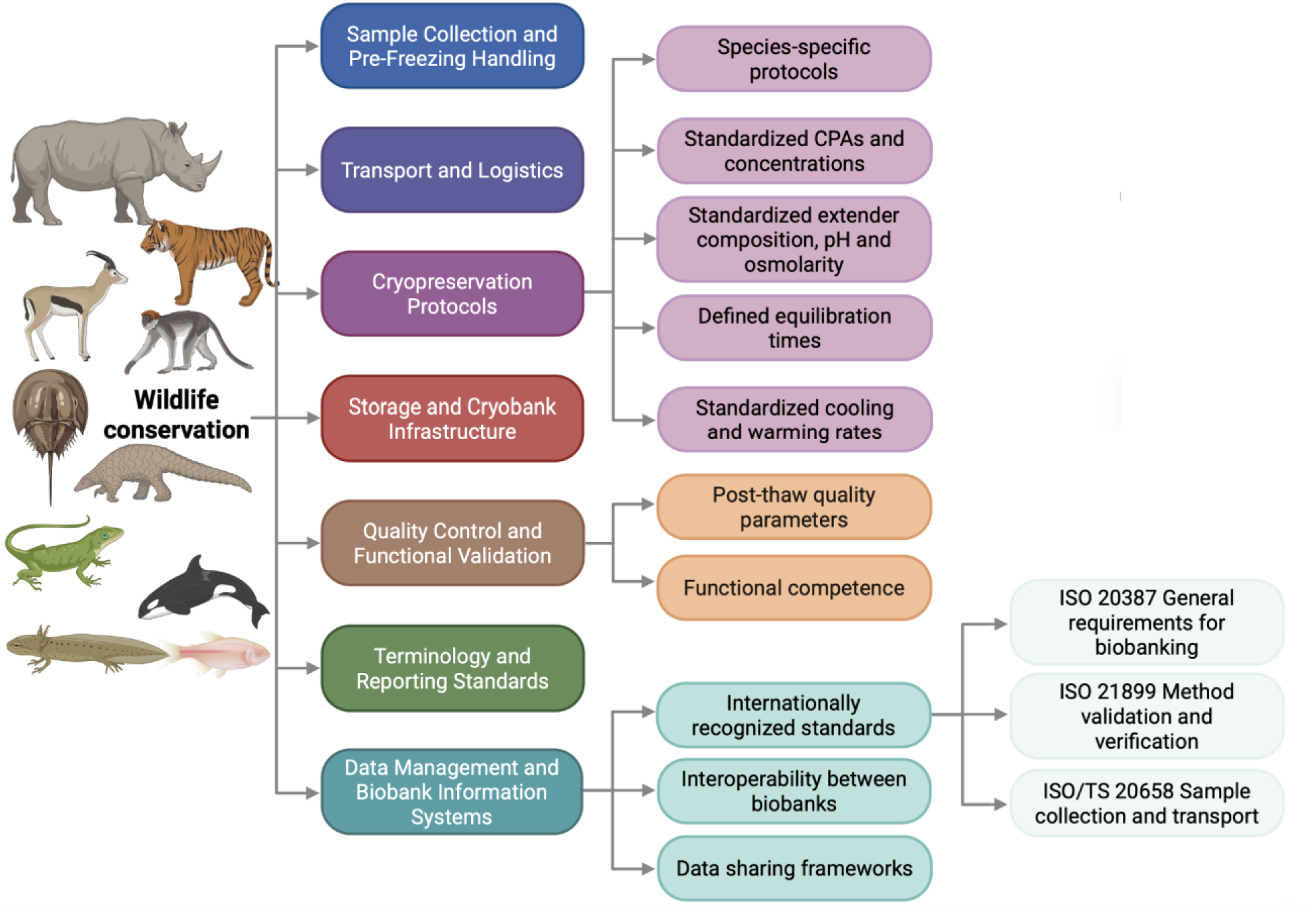
Conceptual framework for the standardization of wildlife cryopreservation and conservation biobanking.

Various levels of regulation must be considered, as well as compliance with international frameworks such as the Convention on Biological Diversity, the Cartagena and Nagoya Protocols ^72^, and the Convention on International Trade in Endangered Species^73^.

The development of effective cryopreservation protocols and their adaptation are complicated by the lack of information on key species-specific characteristics that determine cell cryosensitivity and vary significantly between species and within individuals within a species^11^. This is often complicated, since the selection of species for cryopreservation is largely opportunistic: samples are collected during routine veterinary procedures, capture and release of animals, or necropsies. As a result, the majority of biomaterial comes from zoos and rehabilitation centers, which creates a systematic bias towards already captive and accessible species rather than the most vulnerable wild populations ^69^. Practical limitations—small sample sizes, dangerous working conditions with wild or large animals, limited facilities, and chronic underfunding—slow progress. Postmortem recovery of gametes and gonadal tissues from aging or deceased individuals provides a significant source of valuable genomes, particularly when living animals are acyclic, sterile, or limited in number^74^. An additional complication is the gap between post-thaw survival rates and the actual functional competence of cells. The integration of new tools could transform biobanking by improving tissue survival and reducing reliance on liquid nitrogen and toxic cryoprotectant mixtures (Figure 9).

**Figure 9.**
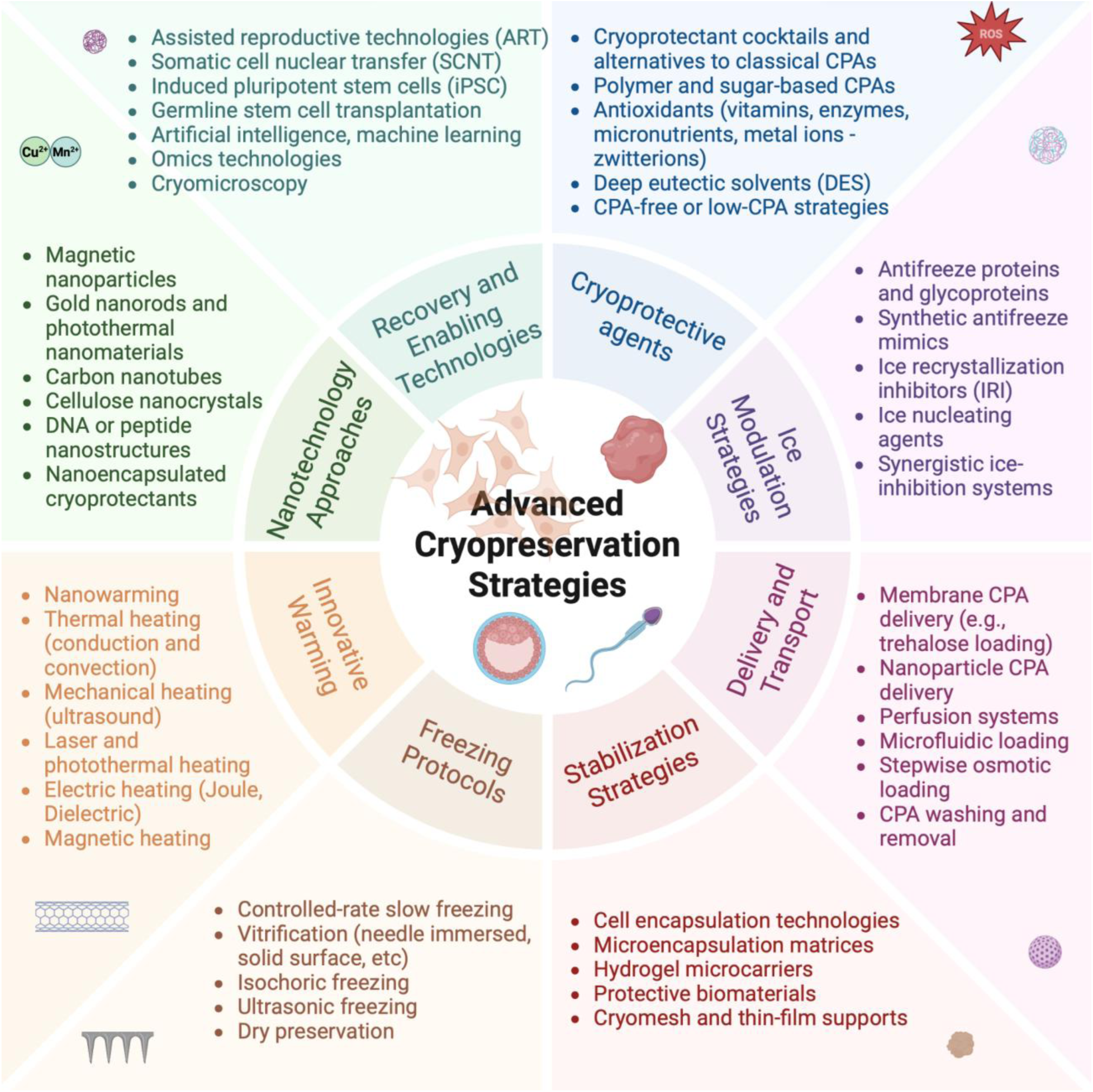
Conceptual framework of advanced cryopreservation strategies for animal cells, tissues, and organs. The inner circle represents eight major strategic domains that includes representative methods or approaches (shown outside the circle) that contribute to enhancing post-thaw viability, structural integrity, and functional preservation. Created with BioRender.

Ultimately, cryopreservation is not an end in itself, but a crucial link in a larger process. Its full value for species conservation is realized only in combination with other technologies, such as ART, embryo transfer, stem cell technologies, the use of iPSCs, and SCNT technology.

The lack of regional networks limits the ability to coordinate sampling strategies, share expertise, and ensure long-term conservation of genetic resources for species whose ranges extend beyond national borders. Given that many Central Asian species, including the saiga antelope (*Saiga tatarica*), snow leopard (*Panthera uncia*), and Asian wild ass (*Equus hemionus*), inhabit transboundary habitats, a transnational biobank for Central Asian biodiversity could unite collections from Kazakhstan, Uzbekistan, Mongolia, and neighboring regions, while providing a common digital infrastructure for cataloging and accessing specimens.

### Conclusions

According to the IUCN Red List, 28% of all assessed species are threatened with extinction, yet developed cryopreservation protocols are lacking for majority of these species. For non-mammalian groups such as amphibians, fish, and corals, reproductive physiology and cryobiology remain largely unexplored, heightening the urgency for fundamental comparative studies using target species and closely related experimental models. In this regard, a stepwise strategy of developing protocols using closely related species and then adapting them to rare species is crucial, reducing the experimental burden on vulnerable populations. Species decline and extinction significantly limit the time window for obtaining biological material and developing effective cryopreservation protocols.

Looking forward, wildlife conservation will require accelerated accumulation of fundamental knowledge about biodiversity and reproductive physiology, the active development of next-generation cryonics tools, and the integration of these technologies into conservation strategies.

### Limitations of the Study

This review and meta-analysis have several limitations that should be considered when interpreting the results. The studies from non-English-speaking countries may have been missed; bias and selective reporting cannot be fully excluded. Because there are no standardized protocols, terminology, or reporting procedures for sharing findings, outcome measures were not uniformly provided. Standard deviations were not consistently included and were therefore derived from reported standard errors, cryoresistance ratios, or extracted from graphical data. Although established statistical transformations and calculations were applied, these procedures introduce an additional layer of uncertainty. These limitations highlight the need for standardized reporting and harmonized methodologies in future cryopreservation studies.

## Materials and Methods

### Search Strategy and Study Selection

This systematic review and meta-analysis were conducted in accordance with the PRISMA (Preferred Reporting Items for Systematic reviews and Meta-Analyses) guidelines. The objective was to evaluate cryopreservation outcomes for wild species, regardless of taxonomic group or tissue type. A literature search was conducted in PubMed, Scopus, Web of Science, and Google Scholar, covering publications from 2020 to 2026. All datasets were imported into Rayyan ^75^, and a single reviewer checked automatically detected duplicates. Two independent, unbiased reviewers screened titles and abstracts according to PICO (Population, Intervention, Comparison, Outcome) meta-analysis criteria. Further screening of the full text and selection of eligible studies was also performed independently by each reviewer. Any disagreements were resolved by discussion until a consensus was reached. The selection process was documented in a PRISMA flow diagram using the Haddaway and co-authors template^76^.

The study included papers that: (a) reported cryopreservation of wild species; (b) reported quantitative results relevant to the survival or function of tissues, cells, or the whole organism; and (c) provided a sufficiently detailed description of the methodology and cryopreservation protocol.

Exclusion criteria included: (a) studies on livestock, common laboratory model organisms, or human material; (b) publications without primary data (e.g., commentaries, editorials); and (c) duplicates, unavailable full text, and languages other than English.

Articles containing data on wild mammal sperm freezing, including sperm characteristics both before and after freezing, were subsequently selected for meta-analysis.

### Assessment of Study Quality

The methodological quality of the studies was assessed using the SYstematic Review Center for Laboratory animal Experimentation’s (SYRCLE) risk of bias tool and the Collaborative Approach to Meta Analysis and Review of Animal Data from Experimental Studies (CAMARADES) quality score checklist ^77–78^. The assessment process also considered the clarity of the experimental design, completeness of reporting, and consistency of outcome measures. Low scores observed in studies are partly explained by field-based or opportunistic wildlife studies, where randomization, allocation concealment, and stable housing conditions are often unfeasible. Bias associated with random assignment of animals was considered low if housing conditions did not differ between groups or if postmortem samples were collected.

### Data Extraction

Data extraction was performed manually using a prepared template. Variables included country, species, population, threat level, sample size, tissue or cell types, cryopreservation protocols, cryoprotectant concentrations, extender composition, and outcome measures. For studies without numerical values, data were extracted from plots using WebPlotDigitizer (https://automeris.io/wpd/). When only standard errors were provided, standard deviations were calculated using the formula SD = SE × √n ^79^.

The IUCN Red List of Threatened Species (https://www.iucnredlist.org) was used to evaluate the global risk of extinction. The species are divided into nine categories: Not Evaluated, Data Deficient, Least Concern, Near Threatened, Vulnerable, Endangered, Critically Endangered, Extinct in the Wild, and Extinct.

The terminology surrounding sperm freezing methods can be ambiguous, with “conventional freezing,” “slow freezing,” and “freezing in liquid nitrogen vapor” terms often used interchangeably. Similarly, direct immersion in liquid nitrogen can be referred to as “ultra-rapid freezing”, “pellet freezing”, and “droplet vitrification”. In this context, we have categorized the methods as follows: standard sperm freezing in liquid nitrogen vapor for 5-15 minutes with prior equilibration is referred to as vapor-freezе (rapid freezing); method that employ controlled temperature programs, specialized chambers, and low cooling rates is referred to as slow-freeze (slow programmable freezing); and the process of solidifying small droplets of a sperm and cryoprotectant mixture by immersing them in liquid nitrogen is classified as pellet-freeze (ultra-rapid freezing in pellets). Additionally, the terms "extender" and "diluent" are used interchangeably, both of which may or may not contain a cryoprotectant, and the concentration may be given relative to the extender or its final dilution.

Most studies were comparative, examining different cryopreservation methods, cryoprotectants, extenders, equilibration times, thawing temperatures, and other protocol variations, resulting in significant within-study variability in post-thaw results. In this meta-analysis, we restricted the dataset to the most relevant experiment for each species to avoid disproportionate weighting of studies reporting multiple experimental variants. Experiments and studies with small sample sizes or thawing of only one sample were also excluded.

### Meta-Analysis

Meta-analysis was performed using the *meta*, *metafor, metacont,* and *dmetar* packages in RStudio 2025.09.2+418 with R version 4.5.1 (Integrated Development Environment for R. RStudio, PBC, USA). Effect size was calculated as the mean difference (MD) between the pre- and post-thaw groups, as continuous outcomes were reported on the same measurement scale. The standardized mean difference (SMD) was also avoided, as it is biased in small samples and may overestimate effects across multiple included studies. Between-study heterogeneity was assessed using I², where values above 75% represent substantial heterogeneity, and tau-squared (τ²), where a large value indicates considerable dispersion of true effects. Following current recommendations by Veroniki and co-authors^80^, the restricted maximum likelihood (REML) method was used to estimate between-study variance for continuous outcomes. Because studies differ in methodology and population, to account for between-study heterogeneity, a random-effects model was employed. To ensure more reliable confidence intervals, the Hartung-Knapp (HK) adjustment was applied, which is more conservative and appropriate for high heterogeneity and a small number of studies. It should be noted that confidence intervals are usually wider with the HK adjustment.

### Sensitivity and Subgroup Analyses

Statistical outliers—studies whose 95% CI shows no overlap with the 95% CI of the pooled effect—were removed, and the effect size was recalculated. To assess the robustness of the pooled estimates, we systematically examined outliers and the influence of individual studies using several diagnostic approaches. A sensitivity analysis was subsequently conducted, excluding the flagged studies, to assess the robustness of each parameter’s effect size (Supplementary Table 6).

For the sensitivity analysis, we first used Baujat plots to identify studies that contributed disproportionately to between-study heterogeneity or had a significant effect on the pooled effect size. Second, influence diagnostics (studentized residuals, Cook’s distance, and DFFITS) were examined. Third, we conducted a leave-one-out analysis. Finally, diagnostics of influential studies were conducted using the Graphic Display of Study Heterogeneity (GOSH) framework, which combines K-means, DBSCAN, and Gaussian mixture clustering algorithms to identify studies associated with atypical model configurations. Subgroup analyses were conducted to explore potential sources of heterogeneity further. Subgroups represented by fewer than three studies were excluded or, where possible, combined into the “Other” category. Differences between subgroups were assessed using random-effects models (Figure 6, Supplementary Table 7).

### Publication Bias

The visual analysis of funnel plots indicated no evidence of publication bias (Supplementary Fig. S25-27). Egger’s regression test also showed no signs of funnel plot asymmetry (all *p* > 0.05, Supplementary Table 7). The trim-and-fill method did not detect any missing studies in any parameters except AI, which had three potentially missing studies on the left side of the funnel plot. The estimate of the corrected effect remained close to the original pooled effect.

### Data Availability

All data analysis supporting the conclusions of this study is available without undue reservation, and the raw data used during the current study will be available from the corresponding author on reasonable request.

### CRediT authorship contribution

GI: writing—original draft, validation, methodology, investigation, formal analysis, data curation, and conceptualization. AH: data curation and investigation. JG: writing—review & editing, supervision. NB: conceptualization,supervision, project administration, methodology, investigation, funding acquisition, writing—review & editing,

### Declaration of Generative AI and AI-assisted technologies in the writing process

The authors declare that they did not use AI technologies in the writing process.

### Funding

This research was funded by the Ministry of Sciences and High Education, Kazakhstan, grant AP26104995 and Nazarbayev University FDCRGP grant #SSH2024005 to N.S.B.

